# Epithelial NSD2 maintains FMOs-mediated taurine biosynthesis to prevent intestinal barrier disruption

**DOI:** 10.1101/2024.05.09.593261

**Authors:** Yue Xu, Chunxiao Ma, Ziyi Wang, Wenxin Feng, Hanyu Rao, Wei Zhang, Ningyuan Liu, Rebiguli Aji, Xiangjun Meng, Wei-Qiang Gao, Li Li

## Abstract

Inflammatory bowel disease (IBD) poses a significant challenge due to its intricate pathogenesis. NSD2, a histone methyltransferase responsible for dimethylating histone 3 at lysine 36, is associated with transcriptional activation. However, the precise role of NSD2 in IBD remains unexplored. In this study, we discovered a downregulation of NSD2 in both the intestinal epithelial cells (IECs) of patients and the IBD mouse model. Deficiency of NSD2 in mouse IECs aggravated epithelial barrier disruption and inflammatory response in IBD. Mechanically, NSD2 loss downregulated H3K36me2 and FMO (taurine-synthesis enzyme) mRNA in IECs, resulting in decreased taurine biosynthesis in IECs. Importantly, supplementation with taurine significantly attenuated the symptoms of NSD2 deficiency-induced IBD. These data demonstrate that NSD2 plays a pivotal role in maintaining FMOs-mediated taurine biosynthesis to prevent intestinal inflammation. Our findings also underscore the importance of NSD2-H3K36me2-mediated taurine biosynthesis in maintaining intestinal mucosal barrier homeostasis.

## 1. Introduction

Inflammatory bowel disease (IBD), which includes Crohn’s disease (CD) and ulcerative colitis (UC), represents a group of chronic inflammatory diseases affecting the gastrointestinal tract, distinguished by persistent immune activation, mucosal inflammation, and tissue damage^1^. The intact epithelium and mucosal barrier collaborate to maintain the integrity of the intestinal barrier and colonic homeostasis. The breakdown of the epithelial barrier leads to the inappropriate movement of gut luminal contents, commensal microbiota, and pathogenic microbes into the intestinal lamina propria. Epithelial barrier dysfunctions causes intestinal inflammation and mucosal injury, ultimately causing intestinal carcinogenesis that affects millions of people worldwide^2^. Due to the complex interplay of molecules and pathways, the pathogenesis of IBD remain largely unknown.

Mounting evidence indicate a strong role for amino acid metabolism in maintaining intestinal epithelial homeostasis^3^. Taurine, one of the most abundant amino acids, can remodel the intestinal microbiota to enhance infection resistance and bolster antitumor immunity^4,5^ and acts as a key regulator of gut homeostasis. However, whether taurine metabolism regulates IBD is undetermined (poorly understood).

Epigenetic regulation, encompassing DNA methylation, histone modification, non-coding RNA (ncRNA) and chromatin interaction, is critical for controlling gene expression changes in gastrointestinal development and homeostasis^6,7^. Various levels of genetic and epigenetic changes contribute to pathogenesis of gastrointestinal diseases, such as IBD and colorectal cancer (CRC). Recent studies have shown that histone modifications play an important role in the occurrence and development of IBD^7^. The post-translational histone modifications, including acetylation and methylation, have a significant impact on various cell types associated with IBD in both normal physiological conditions and during periods of inflammation^8–14^. These studies demonstrate that targeting histone modifying enzymes and their associated chromatin modifications might be a promising therapeutic strategy for the treatment of gastrointestinal diseases.

NSD2, also known as WHSC1 (Wolf-Hirschhorn syndrome candidate gene-1) or MMSET (MM overexpresses multiple myeloma SET domain-containing protein), encodes a histone methyltransferase specific for dimethylation of histone H3 at lysine 36 (H3K36me2)^15^. NSD2 is aberrantly expressed, amplified, or somatically mutated in multiple types of cancers. Through affecting the chromatin accessibility and altering nucleosome structure, NSD2 regulates the transcriptional expression levels of a series of genes and changes their corresponding functions^16–18^. In CRC, NSD2 promotes tumor angiogenesis and leads to a poor prognosis^19^. However, the precise role of NSD2 in the context of intestinal inflammation has not been investigated so far.

In this work, we analyzed the expressions of NSD2 in IBD patients and mouse models and investigated the role of NSD2 during IBD with a conditional Nsd2^Vil-KO^ mouse line and IECs-derived organoids. We demonstrate that NSD2 is reduced in epithelial cells of patients with IBD and show that NSD2-mediated taurine biosynthesis is essential for the formation of proper intestinal barrier function. Deficiency of NSD2 in mouse IECs aggravates epithelial barrier disruption and inflammatory response in IBD. Our findings emphasize that NSD2 plays a crucial role in regulating IEC apoptosis and intestinal inflammation, offering new perspectives on potential therapeutic approaches for IBD treatment.

## 2. Results

### 2.1 NSD2 expression is decreased in IBD

To investigate the role of NSD2 in the pathogenesis of IBD, we examined the expression of NSD2 mRNA in publicly available gene datasets of CD and UC samples. The results indicated that NSD2 mRNA was downregulated in CD and UC specimens compared to those in healthy controls (Fig. 1A). To assess the clinical significance of NSD2 in IBD, we conducted immunohistochemistry (IHC) analyses to evaluate the protein level of NSD2. As shown in Fig. 1B, the expression of NSD2 in IBD patients biopsy specimens was significantly lower than normal colonic epitheliam. Similar to the observations in human IBD, the expression levels of NSD2 and H3K36me2 were also decreased in DSS-induced IBD mice at both the mRNA and protein levels compared to controls (Figures 1C-1F). Taken together, these results suggested a potential link between the NSD2 expression and IBD pathogenesis.

**Figure 1.**
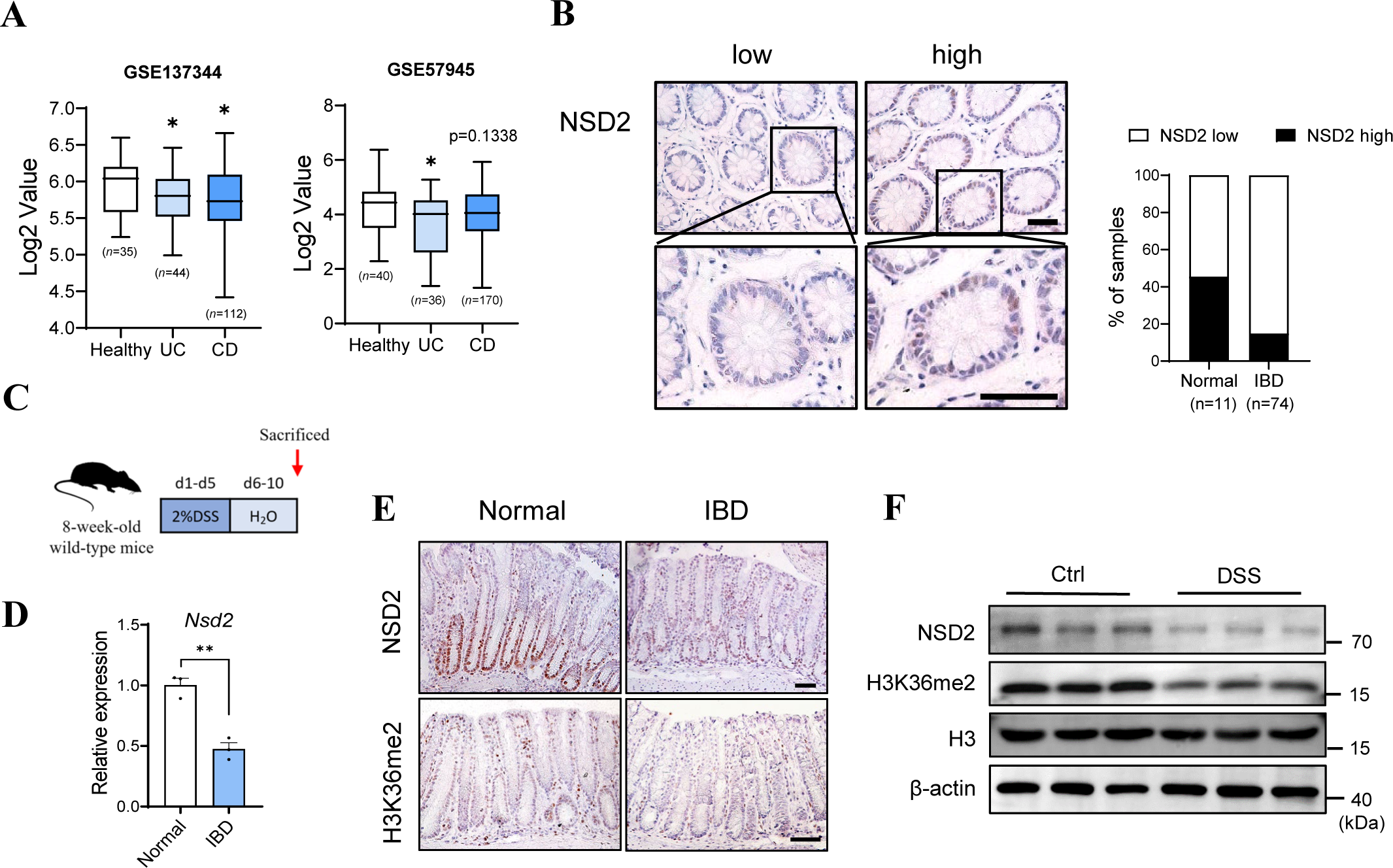
NSD2 expression is decreased in IBD. **A.** Box plot of NSD2 mRNA in healthy controls and UC&CD specimens (using datasets GSE137344 and GSE57945). **B.** NSD2 staining images are shown in the left panel, and epithelial NSD2 expressions in normal and IBD biopsies are quantified in the right panel (χ2 test). Staining indexes use a 10-point quantification scale, and a score > 4 is considered a higher level. **C.** Schematic representation of the DSS protocol used to induce acute colitis. **D-F.** samples are all derived from 8-week-old wild-type mice with or without DSS treatment. **D.** RT-qPCR analysis of NSD2 mRNA in IECs from control and DSS-treated wild-type mice (n = 5 per group). **E-F.** Immunohistochemical **(E)** and immunoblot analyses **(F)** of NSD2 expression in control and DSS-treated wild-type mice are shown. Scale Bars: 50 µm. The data represent the mean ± S.E.M., and statistical significance was determined by a two-tailed Student’s t-test unless otherwise indicated. * p < 0.05, ** p < 0.01, and *** p < 0.001. N.S., Not Significant.

### 2.2 NSD2 deficiency in IECs exacerbates intestinal inflammatory responses

The above results prompted us to use genetically modified mouse models (GEMMs) to investigate the potential function of NSD2 in colon inflammation. Nsd2^flox/flox^ (Nsd2^f/f^) mice and Villin-Cre mice were crossed to generate Nsd2^flox/flox^; Villin-Cre (Nsd2^Vil-KO^) mice. Western blotting and IHC staining results showed that NSD2 was efficiently deleted and the protein level of H3K36me2 was dramatically downregulated in Nsd2^Vil-KO^ mice (Figures 2A-B). Meanwhile, the expression levels of NSD1 and NSD3 remained unchanged (Supplementary Fig. 1a). Nsd2^Vil-KO^ mice exhibited normal intestinal histology, and there were no discernible differences in the number of stem/progenitor cells and terminally differentiated cells between wild-type and Nsd2^Vil-KO^ mice under steady-state conditions (Supplementary Fig. 1b-d). To further investigate the role of NSD2 in IBD, we administered 2% dextran sodium sulfate (DSS) to induce IBD as widely reported in the drinking water of wild-type and Nsd2^Vil-KO^ mice as described^10,20^ (Figures 2C). Nsd2^Vil-KO^ mice exhibited significantly exacerbated IBD compared to control mice, as indicated by accelerated and more pronounced weight loss (Figures 2D), as well as enhanced colon shortening and spleen swelling on day 10 when they were sacrificed (Figures 2E-F). Histopathological examinations revealed that Nsd2^Vil-KO^ mice exhibited more severe phenotypic changes, as evidenced by inflammatory infiltrates, crypt erosion, and an increase in histology score (Fig. 2G).

**Figure 2.**
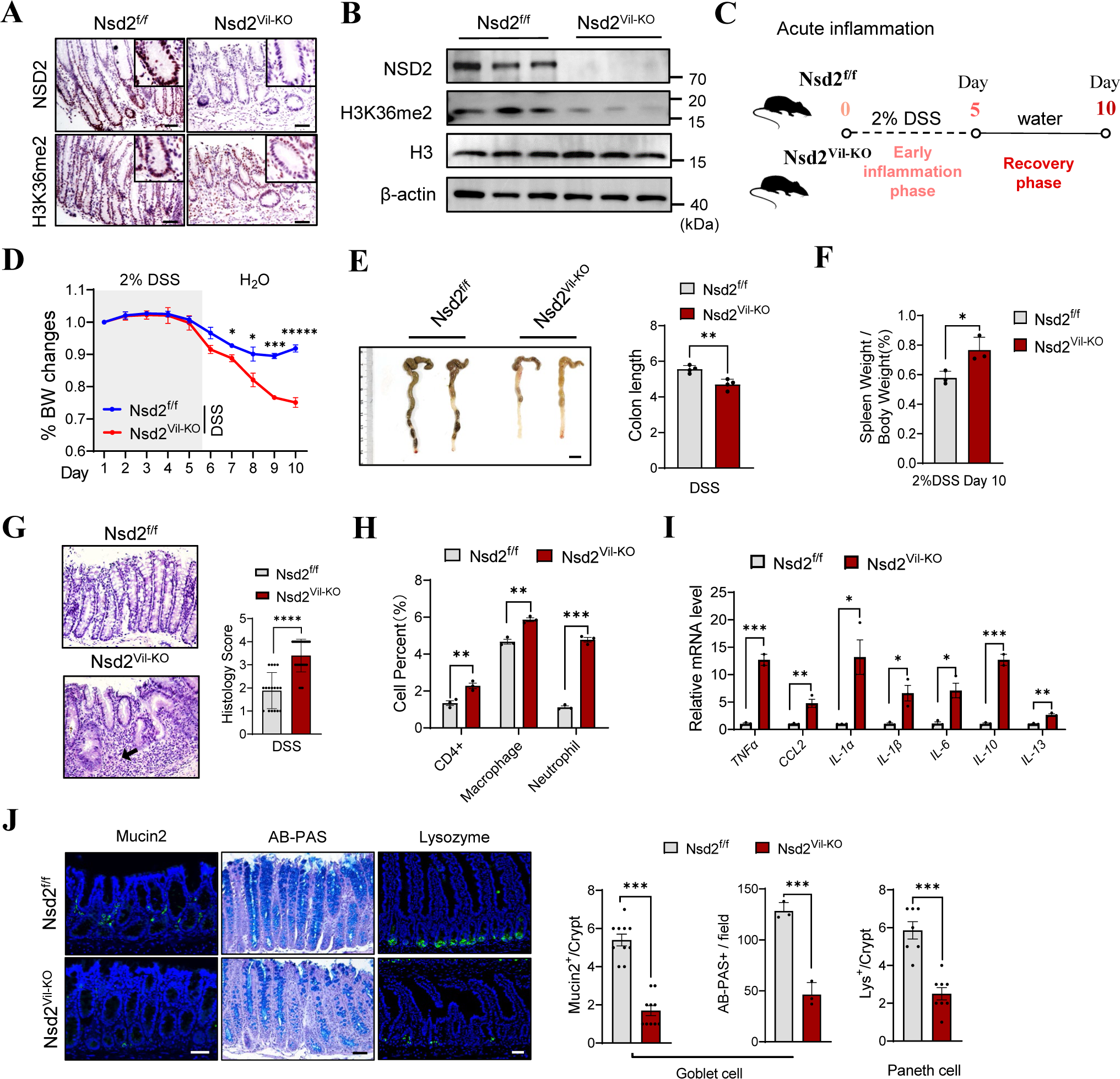
NSD2 deficiency in IECs exacerbates intestinal inflammatory responses. **A-B.** Immuno-histochemical **(A)** and immunoblot **(B)** analyses of NSD2 and H3K36me2 expression in the IECs from Nsd2^f/f^ and Nsd2^Vil-KO^ mice. **C.** Schematic representation of the DSS protocol used to induce acute colitis. **D.** Mice were fed with 2% DSS solution to induce colitis, and weight loss was recorded (n=4). Scale Bars: 1cm. **E.** Representative images of colons from Nsd2^f/f^ and Nsd2^Vil-KO^ mice after DSS treatment (n=3). **F.** Spleen weight normalized to body weight at the time the mice were killed (day 10). **G.** H&E-stained sections and the quantitation of histology score of colon tissue collected on day 10 from 2% DSS-treated Nsd2^f/f^ and Nsd2^Vil-KO^ mice are shown. Scale Bars: 50 μm. **H.** After DSS treatment, colonic lamina propria cells from Nsd2^f/f^ and Nsd2^Vil-KO^ mice are analyzed by flow cytometry (n=3). **I.** Relative mRNA expression levels of inflammatory cytokines and chemokines in the whole colon of Nsd2^f/f^ and Nsd2^Vil-KO^ mice were determined by RT-qPCR (n=3). **J.** Mucin2 (goblet cell), Alcian blue-Periodic acid Schiff (AB-PAS; goblet cell) and lysozyme (Lys; Paneth cell) staining of the intestines from 2% DSS-treated mice and quantitation results are shown on the right (n=3). Scale Bars: 50 μm. All data are presented as mean ± SD, and statistical significance was determined by a two-tailed Student’s t-test unless otherwise indicated. * p < 0.05, ** p < 0.01, and *** p < 0.001. N.S., Not Significant.

We then analyzed immune cell infiltration using flow cytometry. After 5 days of DSS treatment, the numbers of CD4^+^ T cells, macrophages, and neutrophils were increased in Nsd2^Vil-KO^ mice (Fig. 2H). As expected, DSS-treated Nsd2^Vil-KO^ mice exhibited an excessive immune response. RT-qPCR quantitation confirmed a significant increase in the local production of pro-inflammatory cytokines (*Il1a, Il1b, Il6, Il10, Il13, Tnfa, and Ccl2*) in the colon of Nsd2^Vil-KO^ mice (Fig. 2I).

Additionally, NSD2 deletion results in a loss of IECs after DSS treatment (Supplementary Fig. 2a-b). The loss of NSD2 significantly decreased the expression of Mucin2 (Muc2) and Alcian blue–periodic acid–Schiff (AB-PAS), which are produced by goblet cells and form the structure of the intestinal mucus (Figure 2J). Lysozyme staining also showed that Nsd2^Vil-KO^ mice had a reduced number of Paneth cells producing antimicrobial peptides (AMP) in the small intestine compared to the wild-type controls (Figure 2J). Collectively, our results demonstrated that epithelial NSD2 serves as a defense mechanism to prevent DSS-induced IBD.

### 2.3 Epithelial NSD2 deletion exacerbates barrier dysfunction and cell apoptosis

Tight junctions are the primary determinants of barrier function in intact epithelia. Intestinal epithelial damage, such as erosions and ulcers, leads to the loss of tight junctions and therefore causing defects in local barrier function^21^. To investigate the impact of NSD2 on intestinal barrier function in DSS-induced IBD, we examined the expression level and localization of the tight junction protein 1 (ZO-1) and epithelial cadherin protein (E-cadherin) in Nsd2^f/f^ and Nsd2^Vil-KO^ mice. As shown in Figure 3A, Nsd2^Vil-KO^ mice exhibited partially disrupted or discontinuous ZO-1 staining, as well as a substantial reduction in E-cadherin. In addition, Western blot analyses confirmed the reduction of ZO-1, E-cadherin, and Claudin-1 proteins in Nsd2^Vil-KO^ mice (Figure 3B). Furthermore, our results suggested that NSD2 ablation resulted in colonic leakage, as evidenced by the higher serum FITC-dextran concentrations detected in Nsd2^Vil-KO^ mice compared to the controls after DSS treatment (Figure 3C). These findings demonstrated that NSD2 is necessary for maintaining mucosal integrity in an inflammatory environment.

**Figure 3.**
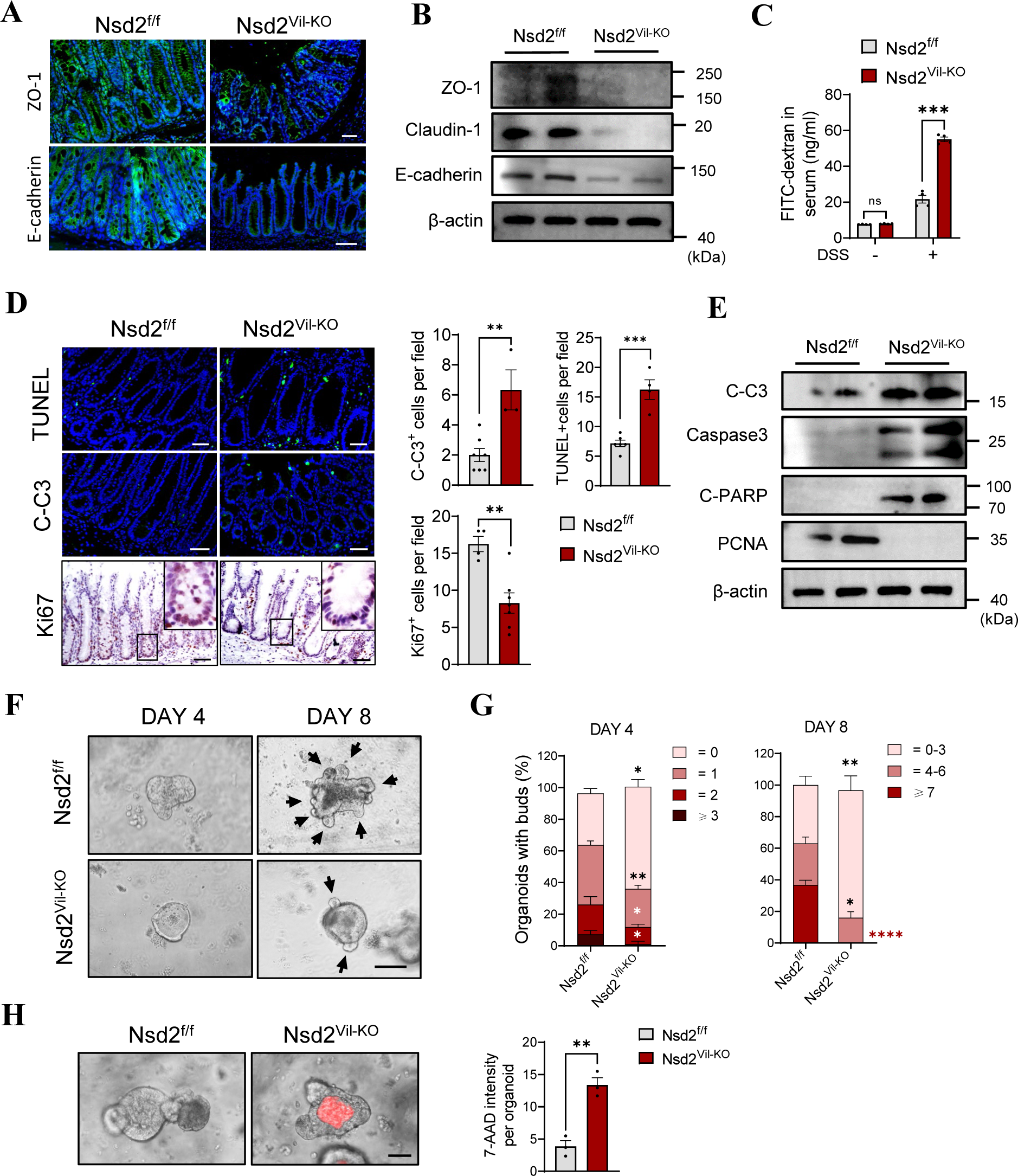
Epithelial NSD2 deletion exacerbated barrier dysfunction and cell apoptosis. **A-B.** Representative immunofluorescence staining **(A)** and immunoblots **(B)** of ZO-1 and E-cadherin in colon sections from DSS-treated Nsd2^f/f^ and Nsd2^Vil-KO^ mice (n=3). Scale Bars: 50 μm. **C.** Colonic permeability was measured by the concentration of FITC-dextran in the blood serum (n=3). **D.** TUNEL (Upper), C-C3 (Medium), and Ki67 (Lower) staining of colon sections and quantitation results are shown (n=3). Scale Bars: 50 μm. **E.** Western blotting analysis of IECs isolated from Nsd2^f/f^ and Nsd2^Vil-KO^ mice. **F.** Organoids derived from the Nsd2^f/f^ and Nsd2^Vil-KO^ mice. **G.** Structure quantification of organoids (n=5, 30 organoids per mouse). Bars represent 100 μm. Quantification was done by measuring the number of the buds. **H.** Apoptotic cells (7-AAD-stained red) in organoids were imaged after 24 h. Quantitation of the fluorescence density per organoid is shown on the right (n=3). The data represent the mean ± S.E.M., and statistical significance was determined by a two-tailed Student’s t-test unless otherwise indicated. * p < 0.05, ** p < 0.01, and *** p < 0.001. N.S., Not Significant.

Intestinal epithelial damage, accompanied by IEC shedding and oligocellular wounds, prompted us to investigate potential defects in cell survival in Nsd2^Vil-KO^ mice^21^. Compared to Nsd2^f/f^ mice, Nsd2^Vil-KO^ mice showed significantly higher numbers of TUNEL- and cleaved caspase-3-positive cells (Figure 3D). Western blot data confirmed the activation of the pro-apoptotic pathway, specifically caspase-3 activation, in colonic crypts from Nsd2^Vil-KO^ mice (Figure 3D). Meanwhile, the number of Ki67-positive cells was decreased in the Nsd2^Vil-KO^ mice (Figure 3D). We also established intestinal organoid cultures to further validate these results. Consistently, loss of NSD2 led to a decrease in size and complexity (higher number of buds; Figure 3E-G) and an increase in cell apoptosis of intestinal organoids (Figure 3H-I).

### 2.4 NSD2 overexpression in IECs prevents DSS-induced IBD

Next, we set out to investigate whether the upregulation of NSD2 could protect the mice from DSS-induced IBD. We generated the mice that overexpress NSD2 specifically in intestinal epithelial cells. Nsd2^OE/+^ mice^22^ were crossed with Villin-Cre mice to produce Villin-Cre; Nsd2^OE/+^ mice (referred to as Nsd2^Vil-OE^ mice) (Fig. 4a). The overexpression of NSD2 in the Nsd2^Vil-OE^ mice was confirmed using IHC and Western blotting (Fig. 4B and C).

**Figure 4.**
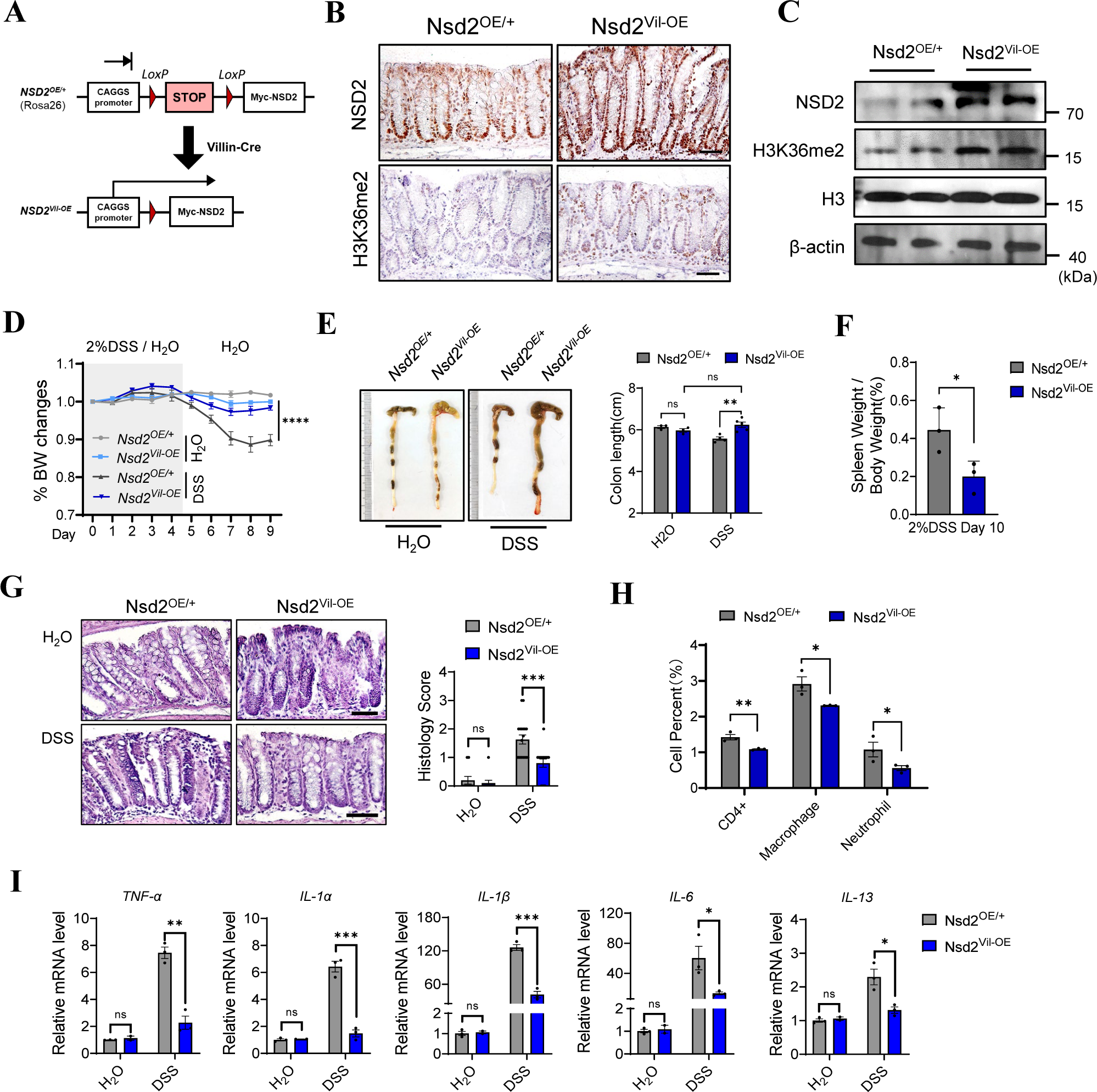
NSD2 overexpression in IECs prevented DSS-induced IBD. **A.** Scheme of Nsd2^OE/+^ mice and conditional overexpression of NSD2 in IECs (Nsd2^Vil-OE^) mice. **B.** Immunofluorescent staining of NSD2 in colon sections from Nsd2^OE/+^ and Nsd2^Vil-OE^ mice. Scale bar, 50 μm. **C.** Immunoblots for NSD2 and H3K36me2 in colonic epithelial cells isolated from colitis mice. **D.** Body weight change of mice treated with or without 2% DSS (n = 5). **E.** The colon length of Nsd2^OE/+^ and Nsd2^Vil-OE^ mice on day 9. **F.** Spleen weight normalized to body weight at the time the mice were killed. **G.** Representative hematoxylin and eosin (H&E) staining images of Nsd2^OE/+^ and Nsd2^Vil-OE^ mice with or without DSS treatment. The histology score is shown on the right. Scale bar, 50 μm. **H.** Colonic lamina propria cells from DSS-treated Nsd2^OE/+^ and Nsd2^Vil-OE^ mice are analyzed by flow cytometry (n=3). **I.** Relative mRNA levels of proinflammatory cytokines in the colon tissue measured by RT-qPCR (n = 3). Data are represented as means ± SD. ns, not significant *P < 0.05, **P < 0.01, and ***P < 0.001.

Nsd2^Vil-OE^ mice and littermates were treated with DSS as described above. By day 10, wild-type mice exhibited IBD syndrome, as indicated by diarrhea, significant body weight loss, and pronounced shortening and thickening of the colon. In contrast, Nsd2^Vil-OE^ mice developed virtually no symptoms, and their large bowel showed few signs of IBD, even compared to the control group that did not receive DSS treatment. (Fig. 4D and E). Histological examination revealed less damage and reduced histological inflammation features in the Nsd2^Vil-OE^ colons, in a sharp contrast to local inflammatory cell infiltration in the wild-type colons (Fig. 4F). Consistently, the colon of Nsd2^Vil-OE^ mice also contained significantly lower levels of proinflammatory cytokines compared to control mice (Fig. 4G). Taken together, our results strongly demonstrated that NSD2 overexpression in IECs is highly protective against experimental IBD in vivo.

### 2.5 Loss of NSD2 exacerbates the inflammatory response in the intestine

To gain an insight into the changes in biological processes and pathways caused by NSD2 loss, we subsequently performed RNA-sequencing (RNA-Seq) analysis using colon epithelial cells isolated from DSS-treated Nsd2^f/f^ and Nsd2^Vil-KO^ mice. The criterion for screening differentially expressed genes was a fold change (FC) > 1.2, and a P-value < 0.05 was considered indicative of significant differential gene expression. Compared to the Nsd2^f/f^ mice, RNA sequencing showed significant changes in the overall transcriptome of the Nsd2^Vil-KO^ mice (Figure 5A). A total of 18,676 differentially expressed genes (DEGs) were identified, including 357 upregulated and 405 downregulated genes (Figure 5B). The gene ontology (GO) analysis for biological processes-associated genes revealed that DEGs were primarily enriched in immune cell migration, response to interleukin-1, and acute inflammatory response, consistent with the earlier results of increased immune cell infiltration (Figure 5C). RT-qPCR was performed and validated these findings (Figure 5D). We conducted Gene Set Enrichment Analysis (GSEA), and the data indicated changes in inflammatory response, consistent with the RNA-seq findings (Figure 5E). These data further confirmed that NSD2 loss exacerbates the inflammatory response in IECs.

**Figure 5.**
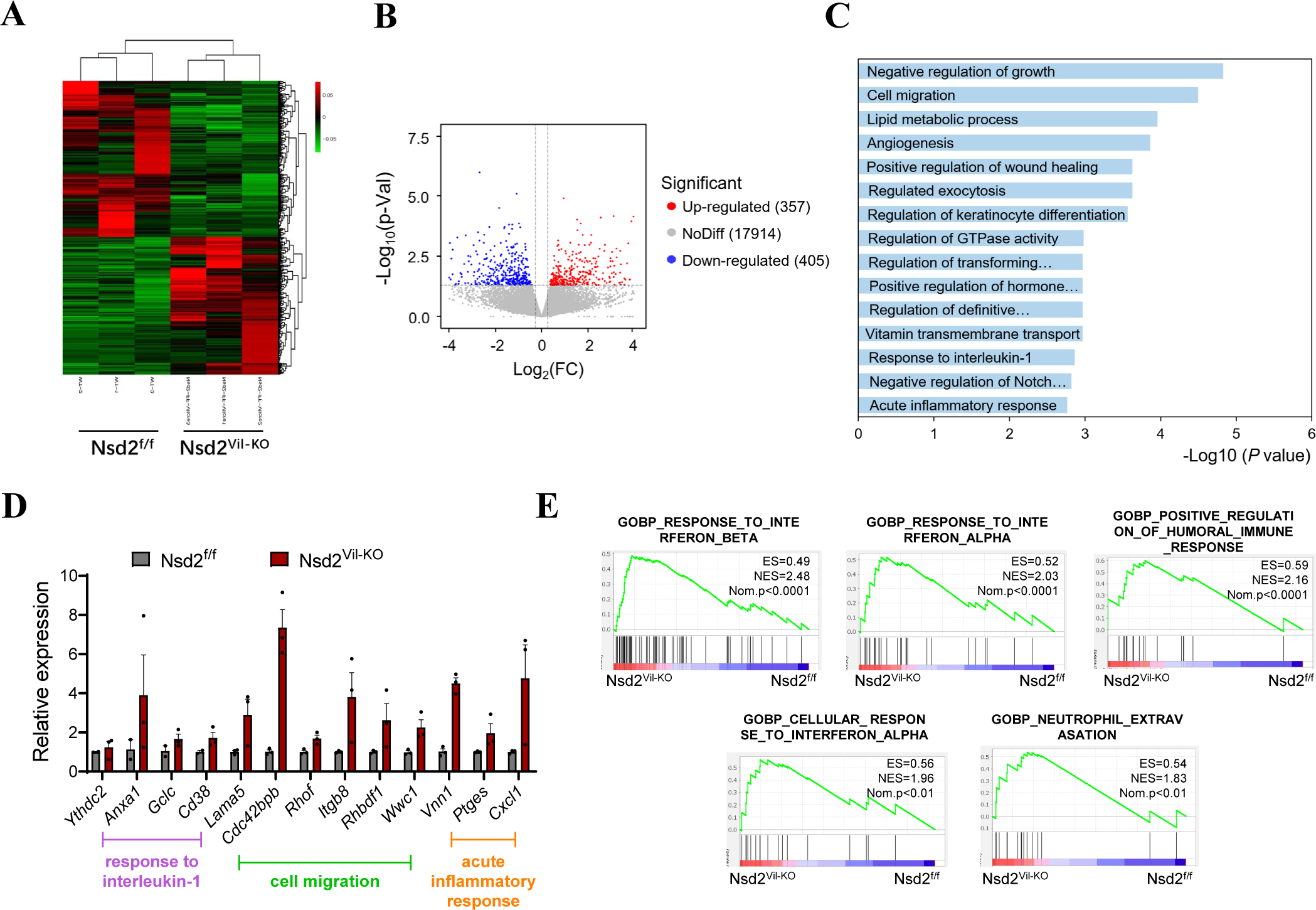
Loss of NSD2 exacerbates the inflammatory response in the intestine. **A.** Heatmap of differentially expressed genes in IECs from 2% DSS-treated Nsd2^f/f^ and Nsd2^Vil-KO^ mice. **B.** Volcano plot of comparative RNA-seq data between 2% DSS-treated Nsd2^f/f^ and Nsd2^Vil-KO^ mice. The x-axis specifies the log_2_ fold changes (FC), and the y-axis specifies the −log_10_ P value. Blue dots represent down-regulated genes, and red dots represent up-regulated genes. **C.** GO analysis of differentially expressed genes. **D.** qRT-PCR was used to verify the differentially expressed genes related to immune cell migration, response to interleukin-1, and acute inflammatory response (n = 3 per group). **E.** GSEA enrichment plots of differentially expressed genes associated with Nsd2 deletion.

### 2.6 NSD2 loss leads to a reduction in Fmo mRNA levels and impedes taurine accumulation

To further explore the NSD2-H3K36me2-mediated mechanism, we performed chromatin immunoprecipitation sequencing (ChIP-seq) targeting H3K36me2 in Nsd2^f/f^ and Nsd2^Vil-KO^ IECs after 4 days of DSS treatment. The H3K36me2 ChIP-seq analysis revealed a total of 10,984 filtered peaks, distributed across promoters (21.11%), introns (35.88%), intergenic regions (40.87%), and exons (2.14%; Figure 6A-B). To correlate the chromatin binding with the transcriptional regulation, the ChIP-seq data were aligned with the expression profile. The Venn diagrams showed that 161 genes had expression changes after NSD2 ablation (Figure 6C). KEGG analysis revealed that most of the differentially expressed genes are related to the FoxO signaling pathway and taurine and hypotaurine metabolism (Figure 6C). Further study showed no significant difference in the FoxO signaling pathway between Nsd2^Vil-KO^ mice and control mice (Supplementary Fig. 3a-c). Notably, the taurine and hypotaurine metabolism pathway was significantly enriched within our dataset. The data showed that *Fmo2*, *Fmo4*, and *Fmo5* (flavin-containing monooxygenase, which functions as the taurine-synthesis enzyme) were downregulated in Nsd2^Vil-KO^ mice. Furthermore, qPCR analysis demonstrated that the loss of NSD2 resulted in the downregulation of FMOs expression levels, while the overexpression of NSD2 led to their upregulation (Fig. 6D). In addition, we conducted qPCR analysis in colorectal cancer cell lines SW620 and MC38 transfected with vector, sg-NSD2, and NSD2-OE plasmids. qPCR analysis revealed that NSD2 deletion significantly downregulated FMOs expression levels, while the expression levels of the FMOs were significantly restored by overexpressing NSD2 (Supplementary Fig. 4a-b). Additionally, ChIP-seq tracks (Fig. 6E) and ChIP-qPCR (Fig. 6F) confirmed that *Fmo2*, *Fmo4*, and *Fmo5* exhibited decreased H3K36me2 occupancy specifically in their promoter regions, compared to controls.

**Figure 6.**
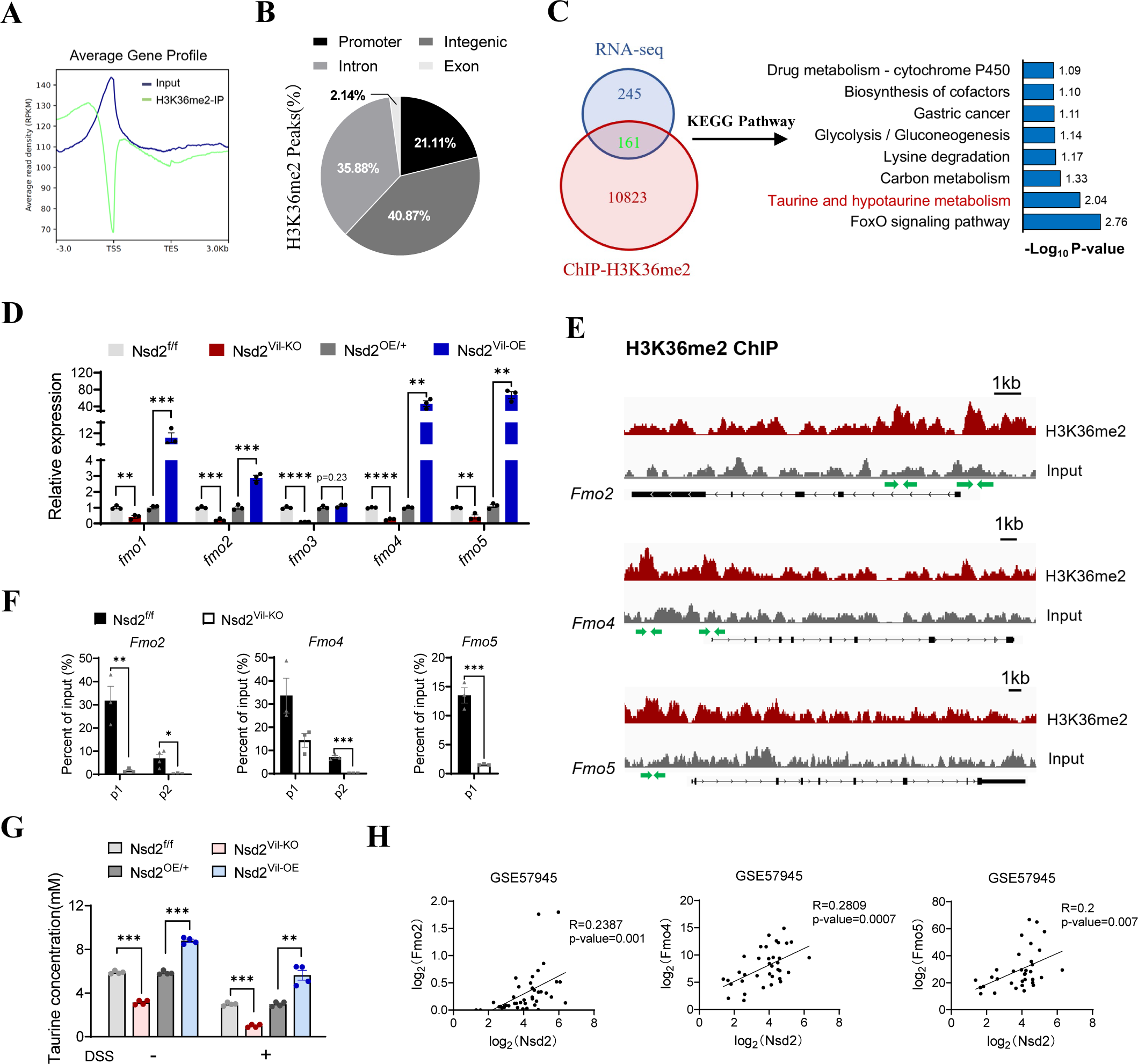
NSD2 loss leads to a reduction in Fmo mRNA levels and impedes taurine accumulation. **A.** Normalized read density of H3K36me2 ChIP-seq signals of colons from Nsd2^f/f^ and Nsd2^Vil-KO^ mice from 3 kb upstream of the TSS to 3 kb downstream of the TES. **B.** Locations of H3K36me2 ChIP peaks relative to genomic annotations. **C.** The overlap of significantly affected genes in ChIP-seq and RNA-seq data. **D.** Representative KEGG terms. **E.** Snapshot of H3K36me2 ChIP-Seq signals at the FMO2, FMO4, and FMO5 gene loci in IECs from DSS-treated (4 d) Nsd2^f/f^ and Nsd2^Vil-KO^ mice. **F.** ChIP-qPCR of FMO2, FMO4, and FMO5. The location of the ChIP primer pairs used is denoted with a green arrow. **G.** The level of taurine in IECs from Nsd2^Vil-KO^ and Nsd2^Vil-OE^ mice and their littermates, treated with or without DSS, were determined using a taurine microplate assay kit (n = 4). **H.** Correlation between NSD2 and FMO2, FMO4, and FMO5 expression levels in IBD specimens (GSE 57945). Statistical significance was determined using the Pearson correlation coefficient. The data represent the mean ± S.E.M., and statistical significance was determined by a two-tailed Student’s t-test unless otherwise indicated. * p < 0.05, ** p < 0.01, and *** p < 0.001. N.S., Not Significant.

Next, we found that the taurine concentration was decreased in Nsd2^KO^ cells compared to the control mice, while the overexpression of NSD2 caused an increase in taurine concentration (Supplementary Fig. 4c). These data indicate a potential inhibition of taurine accumulation in the absence of NSD2. Next, we found a significant reduction in taurine concentration in NSD2-depleted IECs with or without DSS treatment (Fig. 6G). In contrast, a siginificant increase in taurine concentration was seen in Nsd2^Vil-OE^ mice compared to controls (Fig. 6G). Specifically, analysis of clinical IBD patient specimens indicated that there were significantly positive correlations between the mRNA levels of NSD2 and *Fmo2*, *Fmo4*, or *Fmo5*, respectively (Fig. 6H). Taken together, these results coherently demonstrated that NSD2 loss leads to reductions at *Fmo* mRNA levels and impedes physiological taurine metabolism.

### 2.7 The supplementation of taurine relieves intestinal epithelial damage in NSD2-deficient IBD mouse

Given that NSD2 loss in IECs led to a lower FMOs-mediated taurine concentration, we next investigated whether supplementation of taurine could alleviate experimental colitis caused by NSD2 deficiency. We administered 2% DSS in the drinking water of Nsd2^f/f^ and Nsd2^Vil-KO^ mice for 5 days and continued with 5-day taurine supplementation by intragastric administration (Figures 7A). As expected, Nsd2^Vil-KO^ mice showed a significant relief from IBD after supplementation of taurine, as indicated by body weight, colon length, the ratio of splenic weight to body weight and histological score (Figures 7B-E). In addition, the infiltration of immune cells and the induction of proinflammatory cytokines and chemokines were remarkedly inhibited in taurine-supplemented Nsd2^Vil-KO^ mice compared to that in control mice (Figures 7F-H). Furthermore, we observed a recovered protein level of ZO-1 and E-cadherin in distal colon tissues of Nsd2^Vil-KO^ mice with taurine supplementation (Figures 7I-J). These data demonstrated that taurine supplementation alleviates acute colitis and intestinal epithelial barrier damage in Nsd2^Vil-KO^ mice. Moreover, we found that the number of TUNEL- and C-C3-positive IECs was decreased in taurine-treated Nsd2^Vil-KO^ mice compared to controls after DSS treatment (Figures 7K). Taken together, our results demonstrated that supplemented with taurine inhibited IECs apoptosis attenuates the intestinal inflammation caused by NSD2 loss.

**Figure 7.**
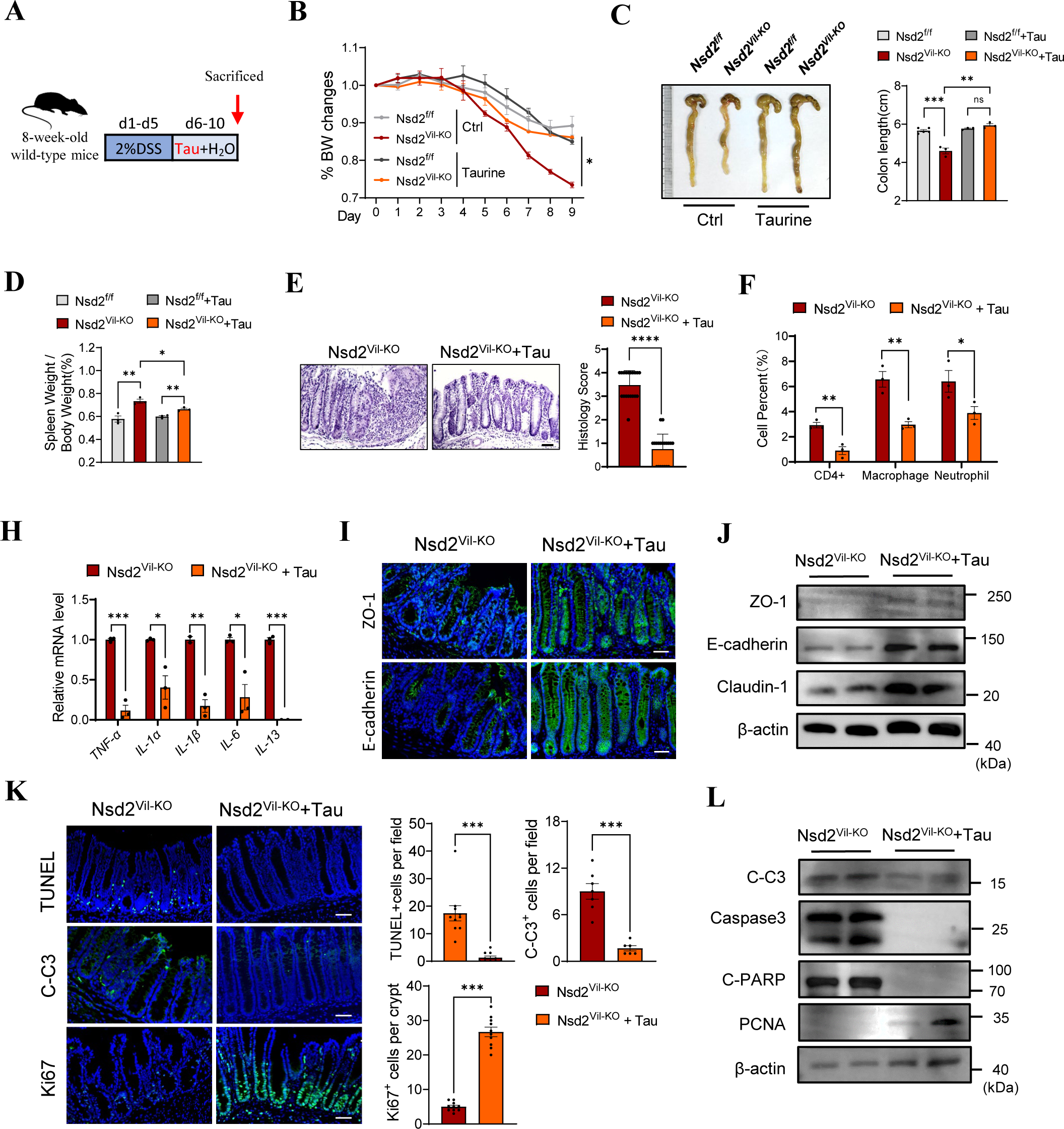
The supplementation of taurine relieves intestinal epithelial damage in NSD2-deficient IBD mouse. **A.** Schematic representation of the DSS treatment and taurine supplementation protocol. Relative percentage change of body weight **(B)**, and representative images and quantification of colon morphology and lengths **(C)** of Nsd2^f/f^ and Nsd2^Vil-KO^ mice after 2% DSS administration for 5 days, intragastric administration for 5 days, and sacrificed on day 11. **D.** Representative images of H&E and histology score (right) of distal colon of Nsd2^f/f^ and Nsd2^Vil-KO^ mice. Scale bar, 50 μm. **F.** Relative gene expression levels of *Tnf-a*, *Il-1a*, *Il-1b*, *Il-6* and *Il-13* from colonic tissues of Nsd2^f/f^ and Nsd2^Vil-KO^ mice. **G-H.** Representative images of ZO-1 and E-cadherin staining of the distal colon from Nsd2^f/f^ and Nsd2^Vil-KO^ mice. Scale bar, 50 μm. **I.** Concentration of FITC-dextran from Nsd2^f/f^ and Nsd2^Vil-KO^ mice was measured to detect intestinal permeability. **J.** TUNEL and C-C3 staining of the distal colon from Nsd2^f/f^ and Nsd2^Vil-KO^ mice and counts of TUNEL- and C-C3-positive cells. Scale bar, 50 μm; n = 3 each group. Data are representative of three independent experiments; error bars show means ± s.d. P values were determined by unpaired two-tailed t-test. For body weight curves, two-way ANOVA analysis with Sidak’s multiple comparisons test.

## 3. Discussion

In this study, we reported that the deletion of NSD2 in IECs leads to intestinal barrier dysfunction and exacerbates inflammatory infiltration. Further analysis revealed that a deficiency of NSD2 results in decreases of H3K36me2 at protein level and Fmo expression at mRNA level to impede taurine accumulation, both in vitro and in vivo. Supplementation of taurine can effectively alleviate intestinal epithelial damage in NSD2-deficient IBD mice. Thus, these results indicate that NSD2 act as a protector of the intestinal epithelial barrier during intestinal inflammation, which helps maintain intestinal epithelial homeostasis by preventing cell apoptosis (Figures 8).

**Figure 8.**
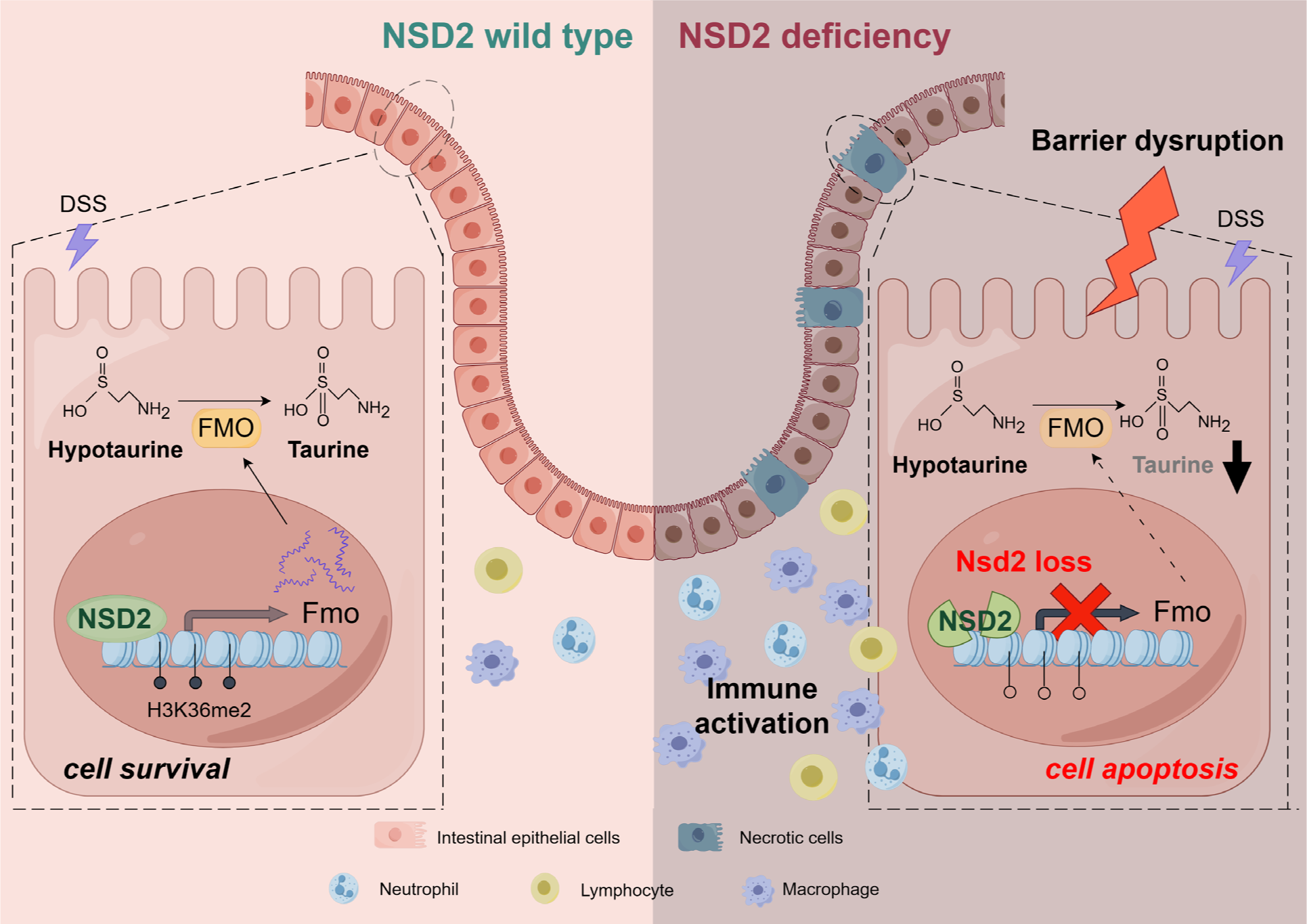
Schematic representation of the mechanism by which NSD2 drives FMO in intestinal taurine accumulation to maintain intestinal homeostasis.

NSD2, a member of nuclear receptor-binding SET domain protein (NSD) family, plays a crucial role in gene transcription, DNA replication, and cellular differentiation. Several studies have shown the significant importance of NSD2 in chromatin regulation and regulates cell senescence, proliferation, migration, invasion, and epithelial-mesenchymal transition (EMT)^19,23–36^. Previous studies suggest that NSD2-mediated H3K36me2 is crucial for transcription activation and the expression of multiple oncogenes in colorectal cancer (CRC)^25,30,37,38^. These studies imply that NSD2, as a histone modifier, may play different roles depending on the genetic environment or context. Till now, there is limited documentation about the role of NSD2 in IBD. In the present study, we report that NSD2 protects the IECs from excessive apoptosis to facilitate the maintenance of intestinal epithelial homeostasis through FMOs-mediated physiological taurine metabolism.

Taurine is a conditionally essential beta-amino acid derived from cysteine in mammals. It is an essential nutrient for the development, growth, and maintenance of physiological functions in various tissues^39^. Low taurine concentration is closely correlated with several age-related diseases^40^ and taurine deficiency causes functional impairments in skeletal muscle, eye, and the central nervous system^41^. Moreover, taurine supplementation could enhance resistance to infection, prevent diabetes mellitus, insulin resistance and its complications, ischemic stroke, and inflammation ^4,42–44^. The latest studies reported that taurine metabolism influences the proliferative, anti-apoptotic, and migratory abilities of cancer cells^5^. Taurine can be obtained from the diet and absorbed by cells through taurine transporters^45^. Due to limited exogenous intake, the maintenance of abundant taurine content in the mammalian body also originates from endogenous synthesis^46^. Previous studies have shown that endogenous taurine is primarily produced from cysteine through the action of cysteine sulfinic acid decarboxylase (CSAD)^47^. In addition to the cysteic acid pathway, hypotaurine can be oxygenated to taurine by flavin-containing monooxygenase (FMOs)^46^, which is currently considered the primary pathway for taurine synthesis^46^. Therefore, FMOs are essential for maintaining endogenous taurine synthesis. FMOs, which include subtypes 1–5, function as the taurine-synthesis enzyme that converts the precursor hypotaurine to taurine^46,48,49^. Additionally, FMOs are promiscuous enzymes that metabolize a wide range of exogenous compounds^50^. There were significant differences in the expression levels of FMOs based on tissue, age, sex, and species. In humans and mice, FMOs are primarily expressed in the liver and skin, and to a lesser extent in other tissues, such as the small intestine^48^. Notably, it is reported that FMO5 play important roles in intestinal inflammation^4,51,52^. Recent studies provide new evidence that the FMO family can be influenced by the transcriptional and posttranscriptional regulation of various factors, including the precursor amino acids of taurine, hormones, bile acids (BAs), and cytokines^46^. However, little is known about the epigenetic mechanism of FMOs regulation. Here, we found that the epigenetic modifier NSD2 can enhance the transcription levels of *Fmo2*, *Fmo4*, and *Fmo5* by marking H3K36me2 on their promoters. Firstly, NSD2 deficiency results in the reductions in *Fmo1-5* mRNA levels and decreased taurine concentration in IECs, leading to epithelial barrier dysfunction during intestinal inflammation. Secondly, overexpression of NSD2 in IECs significantly increased FMOs levels and taurine concentration, effectively mitigating DSS-induced IBD. In vitro experiments also confirmed these results. Finally, taurine supplementation inhibited IEC apoptosis and assisted in mucosal repair, thereby effectively attenuating the intestinal inflammation caused by NSD2 loss.

Previous studies have shown that histone modification modulates the intestinal immune microenvironment. For example, a histone deacetylase inhibitor induces apoptosis and improves the function of regulatory T cells in mice with colitis^53^. Zhou et al. reported that suppressing EZH2 (a major histone methyltransferase of histone 3 lysine 27) activity ameliorates experimental intestinal inflammation by decreasing the MDSC populations^11^. Liu et al. also found that EZH2 promotes the inflammatory response and apoptosis in colitis by modulating the TNFα signaling pathway^14^. In addition, Liu et al. reported that SETD2-mediated H3K36me3 can regulate oxidative stress to alleviate inflammation in mice^13^. It is worth pointing out that the role of H3K36me2 in IBD remains unknown. Here, we found that NSD2-mediated H3K36me2 protects IECs from excessive apoptosis by modulating FMOs transcriptional levels and maintaining taurine concentration, thus preserving an intact and functional mucosal barrier.

In summary, we report that the deficiency of NSD2 in mouse IECs results in a decrease in the expression of FMOs at the mRNA level and hinders taurine accumulation, which promotes IEC apoptosis and exacerbates colitis in mice. Therefore, NSD2 plays a pivotal role in the amino acid metabolism of IBD and may also provide therapeutic insights into understanding other human disorders associated with NSD2 deficiency.

## Methods and materials

### Mice

All mice were maintained in a specific-pathogen-free (SPF) facility, and all experimental procedures were approved by the institutional Biomedical Research Ethics Committee of the Shanghai Institutes for Biological Sciences or the Institute of Zoology, Chinese Academy of Sciences. NSD2-flox mice (Nsd2^f/f^) and NSD2-overexpressing mice (Nsd2^OE/+^) were generated as previously reported^22^ and gifted by Prof. Jun Qin. Villin-Cre mice were purchased from Shanghai Bomade Biotechnology Co. Ltd. These strains were interbred to generate the experimental cohorts, which include the following genotypes: Nsd2^Vil-KO^; Nsd2^Vil-OE^. Mice were harvested at the indicated time for intestinal histology investigation. Mice that lose more than 20% of their weight within one week will be euthanized and recorded as deceased. All the mice were maintained on a C57BL/6J background, and littermates with the same treatment were used for control experiments. For permeability experiments, 2-month-old mice were fasted for 4 hours and then treated with FITC-conjugated dextran (500 mg kg^−1^ body weight). The fluorescence intensity was determined using a FITC-dextran standard curve. To induce colitis, mice at 2 months of age were fed 2% DSS^20^ (molecular weight, 36-50 kDa; MP Biomedicals) for 5 days, followed by regular drinking water. Body weight was recorded daily. To investigate the protective capability of taurine, the mice were exposured to 2% DSS for 5 days, and the taurine supplementation group was given 1000 mg/kg taurine by intragastric administration daily. The control group was given distilled water (n = 3 for each group). At the end of the experimental period, the animals were euthanized, and the intestine were collected for the following experiments.

### Human specimen analysis

Patients with IBD and non-IBD control subjects for this study were recruited from Renji hospital (Shanghai, China). The use of pathological specimens, as well as the review of all pertinent patient records, were approved by the Ethics Review Board of Renji hospital. Immunohistochemical analyses were performed using anti-Nsd2, Abcam, Cat# ab75359 (1:500); anti-H3K36me2, Abcam, Cat# ab9049 (1:1000). Protein expression was scored based on a multiplicative index of the average staining intensity (0–3) and the extent of staining (0–3), which yielded a 10-point staining index that ranged from 0 to 9.

### Taurine measurements

Levels of taurine were detected in the IECs and cell lines by Taurine Microplate Assay Kit (Absin, China) according to the manufacturer’s instructions.

### IEC isolation

IECs were isolated from Nsd2^Vil-KO^, Nsd2^f/f^, Nsd2^Vil-OE^ and Nsd2^OE/+^ mice. IECs were isolated from mice colons. The colon tissue obtained from mice was cut into small fragments of about 1 mm and incubated at 37°C for 15 minutes with 8 mM EDTA. After 15 minutes, replace the EDTA with PBS and shake vigorously for 45 seconds. Repeat the procedure, and combine the supernatant obtained twice. After filtration with a 100-μm cell sieve, the supernatant was centrifuged at 2000 rpm at 4°C for 2 minutes. Pour out the supernatant and resuspend the pellet in cold PBS.

### Serum cytokine quantification

Mouse TNF-α, IL-6, IL-1α, IL-1β, and IFN-γ concentrations in the serum were quantified using ELISA kits purchased from MultiScience, following the manufacturer’s instructions.

### Organoid culture and analysis

The intestines were opened longitudinally, and the villi were scraped away. After thorough washing in cold PBS, the pieces were incubated in 2 mM EDTA/PBS for 10 min at 4°C. Then, the EDTA solution was replaced with PBS and shaken vigorously for 45 s. Crypt fractions were purified through successive centrifugation steps. Hundred microliters of Advanced DMEM/F12 (Invitrogen) containing growth factors (50 ng ml^−1^ EGF, PeproTech; 500 ng ml^−1^ R-spondin, PeproTech; and 100 ng ml^−1^ Noggin, PeproTech) were added and refreshed every 2 or 3 days. After 48 h, the organoids were stained with 7-AAD for 5 min, and photos were captured and quantified using ImageJ software.

### RNA-Seq and ChIP-Seq analysis

IECs were isolated from DSS-treated Nsd2f/f and Nsd2Vil-KO mice using EDTA-based isolation. Each sample contained three to five independently repeated animals and underwent HiSeq sequencing conducted by Guangzhou RiboBio Co., Ltd. Paired-end reads were aligned to the mouse reference genome mm10 using HISAT2. HTSeq version 0.6.0 was used to quantify the reads mapped to each gene. The expression levels of all samples were presented as RPKM (expected number of Reads Per Kilobase of transcript sequence per Million base pairs sequenced), which is the recommended and most common method to estimate the level of gene expression. The statistically significant differentially expressed genes were obtained using a p-value threshold of <0.05 and a fold change ≥1.5. All differentially expressed mRNAs were selected for GO analysis which was conducted using the DAVID database (https://david.ncifcrf.gov/). ChIP-Seq analysis was performed by Active Motif, Inc. using an antibody against H3K36me2 (Abcam). Seventy-five nucleotide reads generated by Illumina sequencing were mapped to the genome using the BWA algorithm with default settings. In total, 26,349 peaks were identified over the input control, and the heat maps and average profile for TSS were generated using ngsplot v2.61.

### Histology, haematoxylin–eosin staining and immunohistochemistry

Tissues were fixed overnight in 4% paraformaldehyde, using the Swiss roll technique, embedded in paraffin, and cut into 7-μm sections. H&E-stained sections were evaluated by a pathologist in a blinded manner. Colitis scores were assigned based on a multiplicative index of the severity of inflammation (0–3), ulceration (0–3), and hyperplasia of the mucosa (0–4)^54^. To quantify goblet cells, PAS+ cells were counted in five random fields from at least five mice per genotype. To quantify Paneth cells, the number of lysozyme-positive cells was counted in 100 crypts from at least 3 mice per genotype. The data are presented as mean ± SEM. The statistical significance between groups was determined using the chi-square test. Primary antibodies used for IHC were as follows: anti-Ki67 (B56), BD Biosciences, Cat# 550609 (1:500); anti-Nsd2, Abcam, Cat# ab75359 (1:500); anti-H3K36me2, Abcam, Cat# ab9049 (1:1000). Biotinylated secondary antibodies were purchased from Jackson Immunology. Staining was visualized with ABC Kit Vectastain Elite (Vector Laboratories) and DAB substrate (Vector Laboratories). PAS staining was performed using PAS staining kits (Sigma-Aldrich) and ALP activity assay was performed using the Alkaline Phosphatase Staining Kit (Vector Laboratories) as described in the manufacturer’s protocols.

### Immunofluorescence

The sections were deparaffinized and rehydrated before antigen retrieval and then cooled to room temperature. Membrane permeabilization was performed using 0.5% Triton X-100, followed by PBS washing and blocking with 10% BSA. The primary antibody was then applied to the samples and incubated at 4°C overnight. DAPI was used for nuclear counterstaining after incubation with the secondary antibody. Primary antibodies used for IF were as follows: anti-Ki67 (B56), BD Biosciences, Cat# 550609 (1:500); anti-ZO-1, invitrogen, Cat# 40-2200 (1:100); anti-E-Cadherin (24E10), Cell Signaling Technology, Cat# 3195T (1:1000); anti-Cleaved Caspase-3 (Asp175), Cell Signaling Technology, Cat# 9661 (1:1000);; anti-lysosome, DAKO, Cat#A0099 (1:500); anti-ChgA, Abcam, Cat# ab715 (1:100); anti-Mucin2 (Ccp58), Santa Cruz Biotechnology, Cat# sc-7314 (1:500); and anti-Lgr5, Abcam, Cat# ab75732 (1:100).

### RNA-seq and ChIP-qPCR assays

Total RNA was extracted from the samples by Trizol reagent (Invitrogen) separately. The RNA quality was checked by Agilent 2200 and kept at −80°C. The RNA with RIN (RNA integrity number) > 7.0 is acceptable for cDNA library construction. First strand cDNA was synthesized using Superscript II (Invitrogen). SYBR Green Master Mix reagents (Roche) and primer mixtures (Supplementary Table 1) were used for the real-time PCR. Student’s t-test was used to statistical analysis and p value <0.05 was considered significant. The ChIP assays were performed using EZ ChIP kit (Millipore). The procedure was as described in the kit provided by the manufacturer. Briefly, isolated IEC cells were fixed by 1% formaldehyde, fragmented by sonication. Anti-H3K36me2, Abcam, Cat# ab9049 (1:1000) was then used for immunoprecipitation. After washing and reverse-crosslinking, the precipitated DNA was amplified by primers and quantified by the qPCR. Primer sequences can be found in the Supplementary Table 1.

### Immunoblotting

The primary antibodies used in this study were as follows: anti-NSD2 (29D1), Abcam, Cat# ab75359 (1:500); anti-H3K36me2, Abcam, Cat# ab9049 (1:1 000); anti-H3, Abcam, Cat# ab10799(1:1000); anti-Caspase3 (8G10), Cell Signaling Technology, Cat# 9665 (1:1000); anti-C-Caspase 3, Cell Signaling Technology, Cat# 9661 (1:1000); anti-PARP (46D11), Cell Signaling Technology, Cat# 9532 (1:1000); anti-ZO1, Invitrogen, Cat# 40-2200 (1:500); anti-PCNA, Santa Cruz Biotechnology, Cat# SC-7909 (1:1000); anti-E-Cadherin, Cell Signaling Technology, Cat# 3195T (1:1000); anti-Claudin1, Cell Signaling Technology, Cat# 4933T (1:1000); anti-Claudin2, Abcam, Cat# ab53032 (1:1000). Raw data of immunoblotting can be found in the source data file.

### FACS

Cell suspensions were subjected to flow cytometry analyses. For the gating strategy, first FSC/SSC was applied to gate the live cells, and then gates were set according to the fluorochrome for subsequent analysis. All the samples in the same experiments and comparisons were gated using the same parameters. The antibodies used in this study were as follows: anti-CD11b-APC, eBioscience, Cat# 47-0112-82 (1:400); anti-F4/80-PE, eBioscience, Cat# 25-4801-82 (1:400); anti-CD4-FITC, eBioscience, Cat# 11-5040-41 (1:400); Gr-1-FITC, eBioscience, Cat# 11-0114-82 (1:400).

### Isolation of lamina propria cells and analysis

Colons were cut into small pieces and then incubated with RPMI medium supplemented with FBS, 0.5 mM DTT, 5 mM EDTA, and antibiotics at 37°C for 30 min. After the removal of the epithelial layer, the remaining colon segments were incubated at 37°C with RPMI medium containing 0.5% collagenase D (Roche) and 0.05% DNase (Roche) for 30 min. Lamina propria cells were stained for surface markers CD4, CD11b, F4/80, and Gr-1 using CD4-FITC, CD11b-APC, F4/80-PE, and Gr-1-FITC antibodies, respectively.

### Statistical analysis

All experiments were performed using 3–15 mice or at least three independent repeated experiments. Unless otherwise indicated, data presented as the mean ± S.E.M. and statistical significance was determined by a two-tailed Student’s t-test. Pearson correlation coefficients were used to evaluate the relationships between NSD2 and gene expressions. χ2 test were used to determine whether there was a significant difference between the expected frequencies and the observed frequencies in one or more categories. *p < 0.05, **p < 0.01, and ***p < 0.001. Reporting summary. Further information on research design is available in the Nature Research Reporting Summary linked to this article.

## Acknowledgments

This work was supported by funds from National Key R&D Program of China (2022YFA1302704 to L.L. and W-Q.G., 2023YFC1404101 to W-Q.G.), National Natural Science Foundation of China (U23A20441 to W.-Q.G., 82372604, 82073104 to L.L.), Science and Technology Commission of Shanghai Municipality (21JC1404100), the Peak Disciplines (Type IV) of Institutions of Higher Learning in Shanghai, 111 Project (B21024) and KC Wong foundation to W-Q.G. and Interdisciplinary Program of Shanghai Jiao Tong University (YG2024ZD11).

## Author Contribution Statement

L.L., W.-Q.G. conceived the experimental concept, designed the experiments, and interpreted the data; Y.X. performed most of the experiments; C.M., Z.W., W.F., W.Z., N.L., R.A., assisted in some experiments; W.-Q.G. assisted in some discussion; Y.X. wrote the manuscript; L.L., provided the overall guidance. All authors read and approved the final manuscript.

## Data Availability Statement

The authors declare that all data supporting the findings in this study are available within the paper, Supplementary information and Source data. All data are available from the authors upon reasonable request.

## Figure legends

**Supplyment figure 1. NSD2 deletion does not affect the self-renewal and differentiation of IECs under steady state.**

**a.** The relative mRNA levels of *Nsd1*, *Nsd2*, and *Nsd3*.

**b.** The relative mRNA levels of intestinal epithelial cell subpopulation markers, including *Sox9, Bmi1, Lgr5, Lyz, Muc2,* and *ChgA* in IECs from 8-week-old Nsd2^f/f^ and Nsd2^Vil-KO^ mice, with n=3 per group.

**c-d.** Representative immunofluorescence staining and quantification of lysozyme (Lys; paneth cell), Mucin2 (goblet cell), ChgA (enteroendocrine cell), and Lgr5 (intestinal stem cell) in IECs from 8-week-old Nsd2^f/f^ and Nsd2^Vil-KO^ mice were shown, with n=3 per group. Scale Bars: 50 µm. The data represent the mean ± S.E.M., and statistical significance was determined by a two-tailed Student’s t-test unless otherwise indicated.

**Supplyment figure 2. NSD2 deletion results in a loss of IECs after DSS treatment.**

**a.** The relative mRNA levels of intestinal epithelial cell subpopulation markers, including *Lyz, Muc2, ChgA, Bmi1,* and *Lgr5,* in IECs from DSS-treated Nsd2^f/f^ and Nsd2^Vil-KO^ mice, with n=3 per group.

**b.** Representative immunofluorescence staining and quantification of ChgA (enteroendocrine cell) in IECs from DSS-treated Nsd2^f/f^ and Nsd2^Vil-KO^ mice were shown, with n=3 per group. Scale Bars: 50 µm. The data represent the mean ± S.E.M., and statistical significance was determined by a two-tailed Student’s t-test unless otherwise indicated. * p < 0.05, and *** p < 0.001. N.S., Not Significant.

**Supplyment figure 3. Adult NSD2 loss does not affect the FoxO pathway in the colon.**

**a.** Real-time PCR analysis was conducted to validate 6 candidate target genes of the FoxO pathway from figure 6C in IECs from 2% DSS-treated Nsd2^f/f^ and Nsd2^Vil-KO^ mice, with n=3 per group. The analysis was performed using an unpaired t-test.

**b.** The relative mRNA levels of Foxo1 and Foxo3 in IEC from 2% DSS-treated Nsd2^f/f^ and Nsd2^Vil-KO^ mice mice were analyzed, with three samples per group. **c.** ChIP-qPCR analysis was conducted to assess H3K36me2 binding for Homer1, Pik3r3, Sgk3, Fbxo32, and Tgfbr1 in IECs from DSS-treated (4 d) Nsd2^f/f^ and Nsd2^Vil-KO^ mice mice, with IgG serving as the control.

**d.** IHC analysis was conducted to assess the levels of FOXO1A, p-FOXO1A, FOXO3A, p-FOXO3A, and FOXO4 in colon tissue from 2% DSS-treated Nsd2^f/f^ and Nsd2^Vil-KO^ mice mice, with 3 mice per group. The data represent the mean ± S.E.M., and statistical significance was determined by a two-tailed Student’s t-test unless otherwise indicated. * p < 0.05, ** p < 0.01, and *** p < 0.001. N.S., Not Significant.

**Supplyment figure 4. Loss of NSD2 reduces Fmo RNA levels and impedes taurine accumulation in vitro.**

**a.** RT-qPCR analysis of Fmo genes was conducted using Nsd2^Ctrl^, Nsd2^KO^, Nsd2^EV^, and Nsd2^OE^ cells (derived from SW620), with n=3 per group. The analysis was performed using an unpaired t-test.

**b.** RT-qPCR analysis of Fmo genes was conducted using Nsd2^Ctrl^, Nsd2^KO1^, Nsd2^KO2^, Nsd2^EV^, and Nsd2^OE^ cells (MC38 derived), n=3 per group. Statistical analysis was performed using an unpaired t-test.

**c.** The level of taurine in Nsd2^Ctrl^, Nsd2^KO^, Nsd2^EV^, and Nsd2^OE^ cells (MC38 derived) was determined using a taurine microplate assay kit (n = 4). The data represent the mean ± S.E.M., and statistical significance was determined by a two-tailed Student’s t-test unless otherwise indicated. * p < 0.05, ** p < 0.01, and *** p < 0.001.

