## Supplement figure for "Epithelial NSD2 maintains FMOs-mediated taurine biosynthesis to prevent intestinal barrier disruption"

**Supplyment figure 1.** NSD2 deletion does not affect the self-renew and differentiation of IECs under steady state.

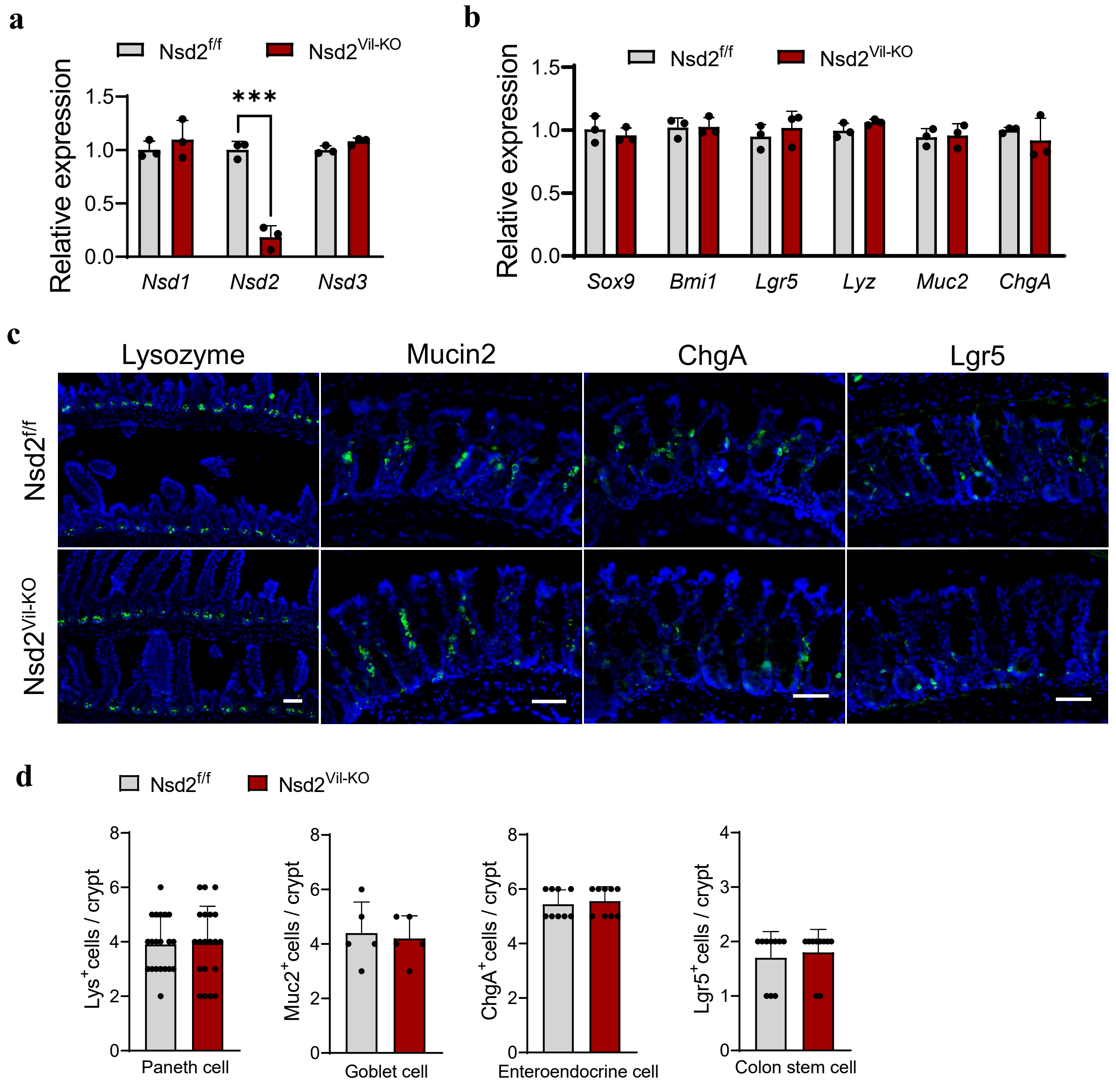

**Supplyment figure 2.** NSD2 deletion results in a loss of IECs after DSS treatment.

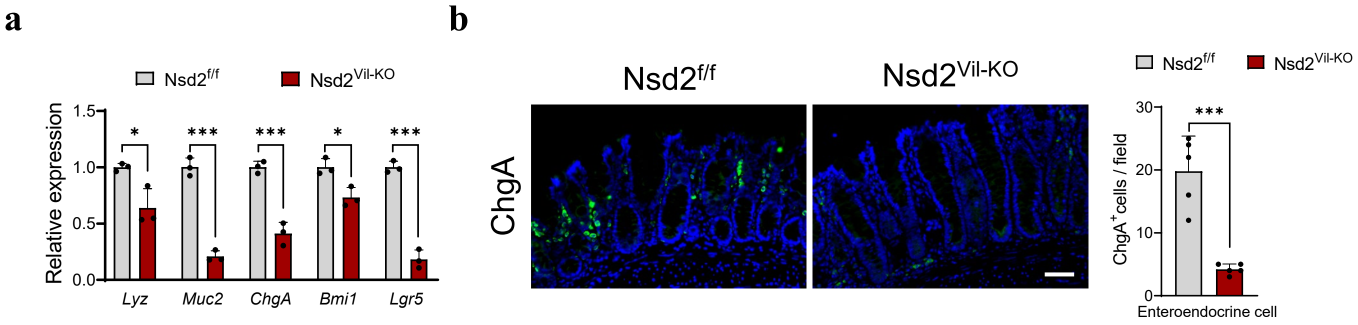

**Supplyment figure 3.** Adult NSD2 loss does not affect FoxO pathway in the colons.

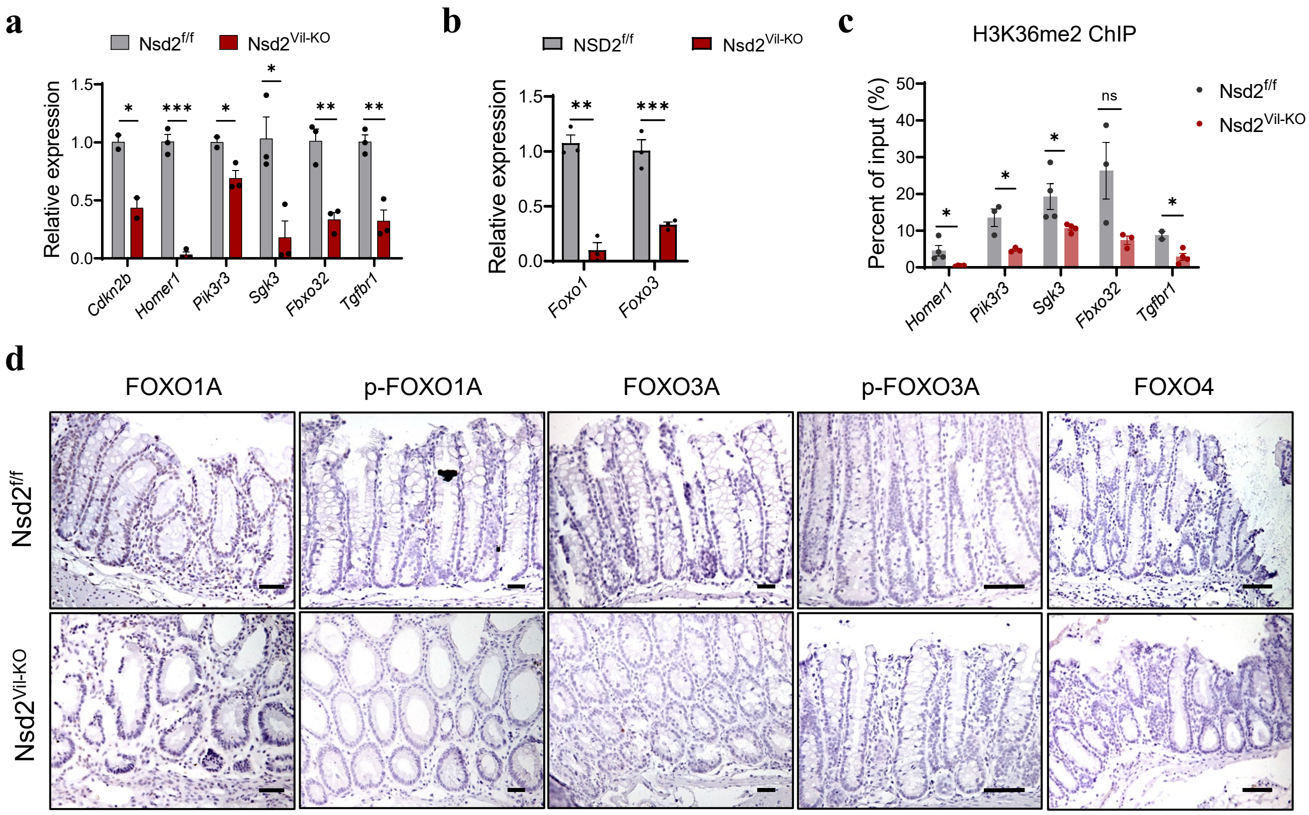

**Supplyment figure 4.** Loss of NSD2 reduces Fmo RNA levels and impedes taurine accumulation in vitro.

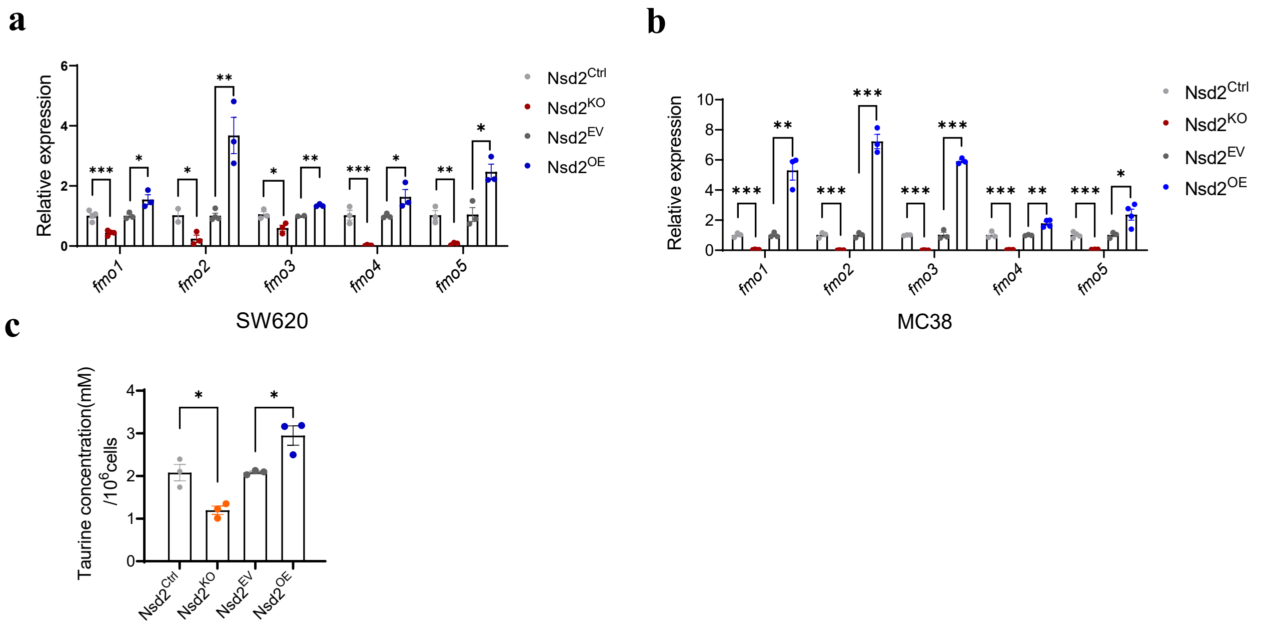
