## Supplement table 1 for "Epithelial NSD2 maintains FMOs-mediated taurine biosynthesis to prevent intestinal barrier disruption"

**Supplemental Table 1: Primers used for qRT-PCR**

|  | Forward primer sequence | Reverse primer sequence |
| --- | --- | --- |
|  | (5’-3’) | (3’-5’) |
| m-GAPDH | AGGTCGGTGTGAACGGATTTG | TGTAGACCATGTAGTTGAGGTCA |
| h-GAPDH | AGAAGGCTGGGGCTCATTTG | AGGGGCCATCCACAGTCTTC |
| β-actin | GATCAAGATCATTGCTCCTCCTG | AGGGTGTAAAACGCAGCTCA |
| h-fmo1 | GCCAAGCGAGTTGCCATTG | CACAGACTTGTAGAGACTGGCT |
| h-fmo2 | GGAGTGTGGAGGTTCAAAGAG | TGCTGGTGTTGGTAACGACAG |
| h-fmo3 | CATCATGTGTATCCCAACCTACC | CTGGTTCTTTATAGTCCCTGCTG |
| h-fmo4 | ATGGGATGACCAGGGTCTATAAG | GTGGAAAGGGAAGTCACTGTAAC |
| h-fmo5 | CCATCACACCAATGCTCATCT | TTCCAGTGAATCCCTCTGGGT |
| ChIP-fmo2-1 | TTAGGTCCTGATGGTGGGGG | GGGAAGCCCAAGGCTTGTAT |
| ChIP-fmo2-2 | AATAAGAGCAAGGGGCAGGG | TGCCTGGCAGAGAGCTAAGA |
| ChIP-fmo2-3 | TCCATACTTGCTGCTTGCCA | TTCACCACCCTTCTGCTGTC |
| ChIP-fmo2-4 | GGGTCAAGGAGGAGGGTCTT | GGCGGTAAGTGACACCATGA |
| ChIP-fmo4-1 | GGGATTCTTCCTGCTGGGAT | TACTGTGCAGAGGAGCACAAG |
| ChIP-fmo4-2 | GTGGTGTGGGTCTTCTGTGA | GACTTCCCCTGGAGGGCTAT |
| ChIP-fmo5-1 | CATCCCAAGGGCCTTCTCAG | GGTTCCAGAGTGGTGTGTGT |
| ChIP-fmo5-2 | CTTCTCCAGCTGGTTTGCCT | AGCCAGAATTTAGCCTCCCG |
| fmo1 | AGTGTGCAGTATAACAAAACGCC | CGTAAAGCAGACTGGGAAGGAAT |
| fmo2 | AGTGGCCTAATCTCTCTGAAGT | CATCGGGAAGTCACTGAAACAG |
| fmo3 | AGCAGGGACTATAAGGAACCAG | GCCTTGCTGTACCACCAGT |
| fmo4 | GATTGGAGCTGGCGTAAGTG | TGTCAGCAAACTTCCACAGTC |
| fmo5 | AGGATTGCTGTAATTGGAGCAG | CCGATGTCACCAGACCTTTCA |
| ChIP-homer1 | TGATAGCAAGCCCTAAAAACATATC | TGCTGAATTGTCTTGGAATGTGA |
| ChIP-sgk3-1 | CCACAGTGGGCTAATGCGAT | TTTTAAAATTGTGTTGGCATTTGGT |
| ChIP-sgk3-2 | CCACAGTGGGCTAATGCGATA | GTTTTAAAATTGTGTTGGCATTTGG |
| ChIP-pi3kr3-1 | ACAAAGGAGTTTGCCACCCA | CAGAGTTGGCCTTGGAGAGG |
| ChIP-pi3kr3-2 | CCACCCAGCCTAATGACCTG | AGTTGGCCTTGGAGAGGTTG |
| ChIP-fbxo32-1 | CTAGGTCAGCAGGCCAAGTC | AAGAGCAGGGGAAATGCCAG |
| ChIP-fbxo32-2 | ACCCGAGCTATCCTCTGACA | TTGGCCTGCTGACCTAGAGA |
| ChIP-tgfbr1-1 | AGCACCACAAATGGTTCCCTC | ATCCGTATTGCATCACCTCCA |
| ChIP-tgfbr1-2 | GCACCACAAATGGTTCCCTCC | TATCCGTATTGCATCACCTCCA |
| foxo1 | CCCAGGCCGGAGTTTAACC | GTTGCTCATAAAGTCGGTGCT |
| foxo3 | CTGGGGGAACCTGTCCTATG | TCATTCTGAACGCGCATGAAG |
| Cdkn2b | CCCTGCCACCCTTACCAGA | CAGATACCTCGCAATGTCACG |
| Homer1 | CCCTCTCTCATGCTAGTTCAGC | GCACAGCGTTTGCTTGACT |
| Pik3r3 | TACAATACGGTGTGGAGTATGGA | GAGTCATTGGCTTAGGTGGCT |
| Sgk3 | AGAAAACAGCCCTATGACAACAC | AGCAACATCTCGGCAGTAAAA |
| Fbxo32 | CAGCTTCGTGAGCGACCTC | GGCAGTCGAGAAGTCCAGTC |
| Tgfbr1 | TCTGCATTGCACTTATGCTGA | AAAGGGCGATCTAGTGATGGA |
| Ccl2 | TTAAAAACCTGGATCGGAACCAA | GCATTAGCTTCAGATTTACGGGT |
| TNFα | CGTCAGCCGATTTGCTATCT | CGGACTCCGCAAAGTCTAAG |
| IL-6 | AGTTGCCTTCTTGGGACTGA | CAGAATTGCCATTGCACAAC |
| IL-1β | GAAATGCCACCTTTTGACAGTG | TGGATGCTCTCATCAGGACAG |
| IL-1α | GAGAGCCGGGTGACAGTATC | TGACAAACTTCTGCCTGACG |
| IL-10 | GCTCTTACTGACTGGCATGAG | CGCAGCTCTAGGAGCATGTG |
| IL-13 | AAGGAGCTTATTGAGGAGCTG | TCAGGGAATCCAGGGCTACA |
| Muc2 | GCCCGTGGAGTCGTACGTGC | TTGGGGCAGAGTGAGGCGGT |
| ChgA | AAGTGCGTCCTGGAAGTCATCTC | GCTTGGCTTTTCTGGCTTGC |
| Lyz | GAGACCGAAGCACCGACTATG | CGGTTTTGACATTGTGTTCGC |
| Lgr5 | CCTACTCGAAGACTTACCCAGT | GCATTGGGGTGAATGATAGCA |
| Bmi1 | ATCCCCACTTAATGTGTGTCCT | CTTGCTGGTCTCCAAGTAACG |
| Nsd2 | GGCCAGAACAAGCTCTTACAA | TGTGGGCTCCCATAAAAGCTC |
| CD38 | TCTCTAGGAAAGCCCAGATCG | GTCCACACCAGGAGTGAGC |
| ANXA1 | ATGTATCCTCGGATGTTGCTGC | TGAGCATTGGTCCTCTTGGTA |
| GCLC | GGGGTGACGAGGTGGAGTA | GTTGGGGTTTGTCCTCTCCC |
| YTHDC2 | GAAGATCGCCGTCAACATCG | GCTCTTTCCGTACTGGTCAAA |
| Lama5 | GCTGGCGGAGATCCCAATC | GTGTGACGTTGACCTCATTGT |
| Cdc42bpb | AAGGTGCGGCTCAAGAAGC | TCCGCCACATACTTGTCGC |
| Rhof | AAGATAGTGATCGTAGGTGACGG | GTACTTCTCAAACACCGACGG |
| Itgb8 | AGTGAACACAATAGATGTGGCTC | TTCCTGATCCACCTGAAACAAAA |
| Rhbdf1 | CTTGCGGAGTGTGAGTATGCC | GGTCTGTGTGATGGATGTCTG |
| Vnn1 | CTTTCCTCGCGGCTGTTTAC | CCTCCAGGTATGGGTAGATCGT |
| Ptges | GGATGCGCTGAAACGTGGA | CAGGAATGAGTACACGAAGCC |
| Cxcl1 | CTGGGATTCACCTCAAGAACATC | CAGGGTCAAGGCAAGCCTC |
